# Evaluating a sepsis prediction machine learning algorithm in the emergency department and intensive care unit: a before and after comparative study

**DOI:** 10.1101/224014

**Authors:** Hoyt Burdick, Eduardo Pino, Denise Gabel-Comeau, Carol Gu, Heidi Huang, Anna Lynn-Palevsky, Ritankar Das

## Abstract

**Introduction:** Sepsis is a major health crisis in US hospitals, and several clinical identification systems have been designed to help care providers with early diagnosis of sepsis. However, many of these systems demonstrate low specificity or sensitivity, which limits their clinical utility. We evaluate the effects of a machine learning algodiagnostic (MLA) sepsis prediction and detection system using a before-and-after clinical study performed at Cabell Huntington Hospital (CHH) in Huntington, West Virginia. Prior to this study, CHH utilized the St. John’s Sepsis Agent (SJSA) as a rules-based sepsis detection system.

**Methods:** The Predictive algoRithm for EValuation and Intervention in SEpsis (PREVISE) study was carried out between July 1, 2017 and August 30, 2017. All patients over the age of 18 who were admitted to the emergency department or intensive care units at CHH were monitored during the study. We assessed pre-implementation baseline metrics during the month of July, 2017, when the SJSA was active. During implementation in the month of August, 2017, SJSA and the MLA concurrently monitored patients for sepsis risk. At the conclusion of the study period, the primary outcome of sepsis-related in-hospital mortality and secondary outcome of sepsis-related hospital length of stay were compared between the two groups.

**Results:** Sepsis-related in-hospital mortality decreased from 3.97% to 2.64%, a 33.5% relative decrease (P = 0.038), and sepsis-related length of stay decreased from 2.99 days in the pre-implementation phase to 2.48 days in the post-implementation phase, a 17.1% relative reduction (P < 0.001).

**Conclusion:** Reductions in patient mortality and length-of-stay were observed with use of a machine learning algorithm for early sepsis detection in the emergency department and intensive care units at Cabell Huntington Hospital, and may present a method for improving patient outcomes.

**Trial Registration:** ClinicalTrials.gov, NCT03235193, retrospectively registered on July 27th 2017.

## Background

Sepsis is a common, life-threatening syndrome that arises from the body’s dysregulated response to infection, and has been declared a global health priority by the World Health Organization [1]. In the United States, sepsis is responsible for a cost of over $20 billion [2] and affects a population of 750,000 annually [3]. Severe sepsis, distinguished by organ failure, may progress to septic shock, presenting with refractory hypotension. An increase in mortality rate from over 10% to near 40% accompanies this escalation in condition severity [4]. In spite of the high prevalence of sepsis syndromes and the associated poor outcomes, the variations in host response and disease progression often inhibit the critical early and accurate diagnosis of sepsis. As demonstrated by the recent proposal of changes to the stages of sepsis (Sepsis-3) [5], there is some controversy in establishing unanimous definitions of clinical sepsis presentations. Yet, numerous studies have reached the consensus that early detection of sepsis and compliance with sepsis treatment bundles can positively impact patient mortality and length of stay [6].

Healthcare systems grapple with accurately identifying sepsis early in disease progression. The increasing availability of data from patients’ electronic health records (EHR) may provide valuable insight into the processes of sepsis disease progression. Existing prospective studies of EHR data-derived tools in clinical settings have been primarily rules-based [7], applying preset score thresholds to classify risk level [8]. However, these studies have often demonstrated subpar sensitivity and specificity [9]. Machine learning algorithms have the potential to improve on rules-based systems through flexibility and learning from patient data, clinical response patterns, and correlative trends. Previous work conducted on sepsis detection machine learning algorithms constructed from EHR data include the retrospective studies of Henry et al. [10], Nachimuthu et al. [11], and Sawyer et al [12].

West Virginia provider Cabell Huntington Hospital (CHH), a 303-bed facility, partnered with Dascena (Hayward, CA) to improve sepsis-related outcomes using a machine learning “algodiagnostic” (MLA). The Dascena MLA was validated for sepsis prediction and detection in several studies [13-15], demonstrating an area under the receiver operator characteristic curve (AUROC) over 0.90 using only six vital signs, in a multicenter cohort study of over 650,000 encounters [16]. In a recent randomized clinical trial, mortality decreased by 12.4 percentage points with use of the MLA, a relative reduction of 58% [17]. Comparison of the MLA to rules-based scores such as Systemic Inflammatory Response Syndrome (SIRS) [18] criteria have shown superior sensitivity and specificity up to four hours in advance of sepsis onset [13]. In this study, we evaluate improvements in CHH sepsis-related in-hospital mortality rate and hospital length of stay with the use of the machine learning algodiagnostic in the emergency and intensive care unit (ICU) patient populations, using a before-and-after study design.

## Methods

### Study Design

This study was designed as a prospective before-and-after study (study registration: ClinicalTrials.gov NCT03235193). Approval for the study was granted by the institutional review board (IRB) at Marshall University. We measured pre-implementation baseline metrics as well as post-implementation metrics in order to determine the effect of the algorithm.

Prior to and during this trial, CHH used Cerner’s St. John’s Sepsis Agent (SJSA). St. John’s issues two types of alerts: 1) a SIRS alert fires when three or more vital signs or lab results fall out of range [18] and 2) a sepsis alert fires when at least two SIRS criteria are met and lab results indicate organ dysfunction. When the criteria are met, an alert appears on an electronic health record (EHR) screen and the provider is notified. Although the criteria are designed to discern sepsis progression, SJSA produces a high false alarm rate due to its low specificity [19]. Low specificity often leads to alarm fatigue, or clinician indifference to the alerts, which results in a delay of treatment during the critical early intervention period [20, 21].

Only CHH’s SJSA was active during the pre-implementation period; during the post-implementation period, both the SJSA and the machine learning sepsis predictor were actively monitoring patients. Pre-implementation data were measured in the emergency department (ED) and intensive care units (ICU) during the period July 1 to July 30, 2017, and post-implementation data were measured in the same units during the period spanning from August 1 to August 30, 2017. All data were collected through CHH’s EHR system, CARE Connect (Cerner Corp, North Kansas City, Missouri).

During the post-implementation phase, all patients over the age of 18 who were admitted to either the ED or ICU were monitored by the MLA for sepsis risk. The MLA assessed each patient for sepsis by extracting real-time data from each patient’s EHR and analysing trends in the patient measurements. Risk prediction scores were computed hourly throughout the duration of each patient’s stay. The MLA used in this study is described in detail in prior prospective [16, 17] and retrospective work [13].

The algodiagnostic was designed to compare trends in each patient’s EHR measurements to confirmed prior sepsis cases in order to accurately detect and predict sepsis. The classifier used to perform the comparison was an ensemble of decision trees. After patient data passed through the classifier, the MLA generated a sepsis risk prediction score between 0 and 100. Healthcare providers were called and informed of a possible sepsis case when a patient’s score exceeded 80. At this point, patients were examined and treated under CHH’s standard sepsis protocol. Patients were monitored by the algorithm for the duration of their stay in the ED or ICU. Additionally, patients continued to be monitored by the SJSA in the post-implementation period. This design ensured that minimal risk was incurred by patients; if the MLA failed to detect a case of sepsis, the SJSA may still have detected sepsis and alerted a clinician.

The primary and secondary outcomes assessed in this study were the sepsis-related in-hospital mortality rate and the sepsis-related hospital length of stay (LOS), respectively, at CHH.

### Data Collection and Analysis

Demographic and clinical information was collected for each patient monitored during the study period. Patients were monitored and clinical data were collected during the duration of their stay in the ED and ICU hospital units, and participants were followed until hospital discharge in order to determine overall sepsis-related in-hospital mortality and LOS. Patients were considered to be “sepsis-related” and included for analysis if they met two or more SIRS criteria at any point during their stay in participating units and were over the age of 18. We classified patients in this manner due to the predictive nature of the MLA. Because the algodiagnostic is designed to identify patients likely to develop sepsis, including only patients who met the 2001 consensus definition criteria could have excluded patients who would have developed sepsis had they not been identified and treated early. The SIRS criteria are closely linked to sepsis diagnostic criteria, and their use in this study ensured that only patients with sepsis or closely related conditions were included in our final analysis.

The MLA determined each patient’s sepsis risk through real-time abstraction of data in the patient EHR. At least one measurement each of systolic blood pressure, diastolic blood pressure, heart rate, temperature, respiratory rate, and peripheral oxygen saturation (SpO_2_) were required for sepsis prediction. Any vital signs not recorded during a given hour were gap-filled using a forward-filling imputation process in which the most recently recorded past measurement was used for sepsis risk score computation. Additionally, although not necessary, the algorithm was able to incorporate lab results such as pH, white blood cell count, and glucose levels when they were available. The MLA monitored each patient hourly and analyzed all clinical measurements as well as hourly changes in measurements, in order to determine sepsis risk.

After the conclusion of the study period, the primary outcome of sepsis-related in-hospital mortality and the secondary outcome of average sepsis-related hospital LOS were calculated. Additionally, we retrospectively compared algodiagnostic performance on patient data from the study to the performance of the SIRS, Modified Early Warning Score (MEWS) [22], and the Quick Sequential Organ Failure Assessment (qSOFA) score [4] on the same data set. No interim analyses were performed before the conclusion of the trial.

Two-sample *t*-tests were used to determine if there was a statistically significant difference in means between the pre-and post-implementation periods for sepsis-related LOS. We used the two-proportion (risk difference) and relative risk (risk ratio) *z*-tests to determine if there was a statistically significant decrease in the in-hospital mortality rate with the use of the algodiagnostic. All tests were single tailed with an alpha level of .05, and were performed using the MATLAB software (R2016a version) developed by MathWorks, Inc. (Natick, MA).

## Results

### Patient Characteristics

Our final analysis included a total of 2,296 sepsis-related cases, which included 1,160 patients in the pre-implementation phase and 1,136 patients in the post-implementation phase. Patient demographics for each period are displayed in Table 1. There were no significant differences in demographics between the two periods. Because all patients were tracked throughout the duration of their hospital stay, no patients were lost to follow-up.

**Table 1.** Demographic characteristics for patients monitored during the study period, based on data abstracted from the EHR.

| **Demographic Overview** | **Characteristic** | **Pre-implementation (%)** | **Post-implementation (%)** |
| --- | --- | --- | --- |
| **Gender** | Female | 43.33 | 39.20 |
|  | Male | 56.67 | 60.80 |
| **Age** | 18-29 | 20.47 | 27.08 |
|  | 30-39 | 16.74 | 13.54 |
|  | 40-49 | 13.96 | 12.55 |
|  | 50-59 | 16.74 | 13.12 |
|  | 60-69 | 15.35 | 21.01 |
|  | 70+ | 16.74 | 12.69 |
| **Ethnicity** | Asian or Pacific Islander | 0 | 0.13 |
|  | Black | 12.45 | 4.88 |
|  | White | 86.27 | 94.37 |
|  | Native Hawaiian | 0.43 | 0 |
|  | Unknown | 0.86 | 0.63 |

### Outcomes

We evaluated the number of sepsis-related in-hospital mortality cases and the mean length of stay for sepsis-related patients before and after MLA integration. The pre-implementation baseline mortality rate was 46/1160 (3.97%, standard error (SE) 0.57%). After MLA implementation, the mortality rate was 30/1136 (2.64%, SE 0.48%) representing a 33.5% reduction (P=0.038). During the baseline period, average sepsis-related length of stay was 2.99 days (SE 0.028); post MLA implementation, sepsis-related length of stay was 2.48 days (SE 0.051), a 17.1% reduction (P<0.001). These results are shown in Figure 1.

**Figure 1.**
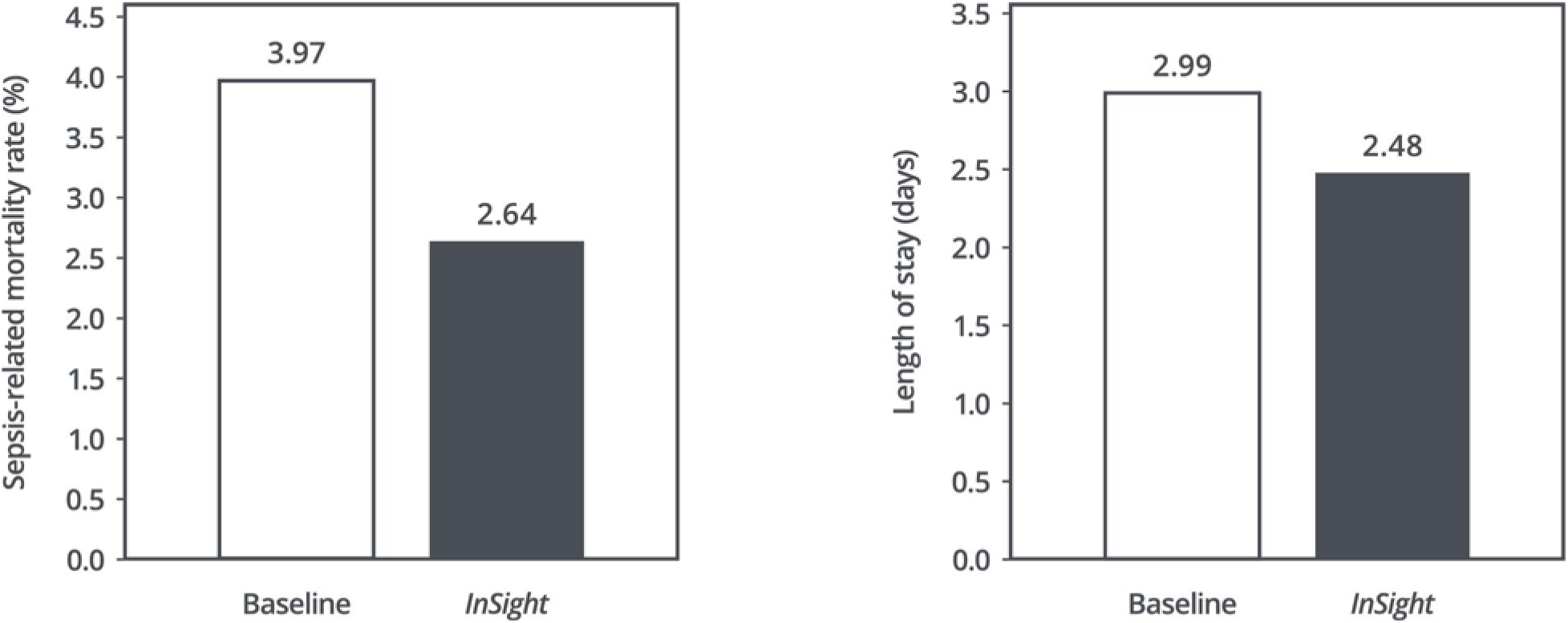
Left: Sepsis-related in-hospital mortality during the baseline period and after machine learning algodiagnostic (MLA) implementation. The use of the MLA was associated with a reduction of in-hospital mortality of 33.5%. Right: Sepsis-related hospital length of stay during the baseline period and after MLA implementation. The use of the MLA was associated with a reduction in length of stay of 17.1%.

In addition to analyzing patient outcomes, we compared the performance of the algodiagnostic to that of the SIRS, MEWS, and qSOFA scores for sepsis detection. On a retrospective set of 1,912 patients (70 meeting severe sepsis criteria) admitted to CHH during the pre-implementation period, the MLA demonstrated statistically higher Area Under the Receiver Operating Characteristic (AUROC) curve, sensitivity, and specificity as compared to all three rules-based scores, with p < 0.05 (Table 2).

**Table 2.**
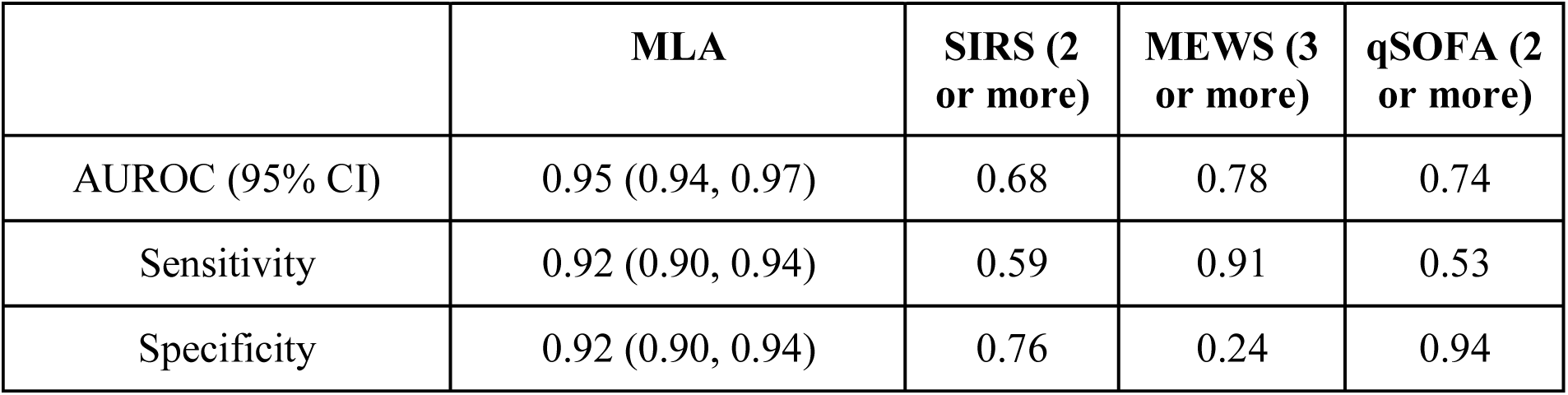
Comparison of area under the receiver operator characteristic curve (AUROC), sensitivity, and specificity for the machine learning algodiagnonstic (MLA), Systemic Inflammatory Response Syndrome (SIRS) score, Modified Early Warning Score (MEWS), and Quick Sequential Organ Failure Assessment (qSOFA) score for sepsis detection. 95% Confidence Intervals (CI) were calculated only for the MLA.

|  | **MLA** | **SIRS (2 or more)** | **MEWS (3 or more)** | **qSOFA (2 or more)** |
| --- | --- | --- | --- | --- |
| AUROC (95% CI) | 0.95 (0.94, 0.97) | 0.68 | 0.78 | 0.74 |
| Sensitivity | 0.92 (0.90, 0.94) | 0.59 | 0.91 | 0.53 |
| Specificity | 0.92 (0.90, 0.94) | 0.76 | 0.24 | 0.94 |

## Discussion

In this prospective study, sepsis-related patient outcomes were improved through the implementation of a machine-learning based sepsis prediction algorithm. When deployed in the emergency department and intensive care units at CHH, the machine learning algodiagnostic resulted in decreases of sepsis-related in-hospital mortality and sepsis-related length of stay (Figure 1). Mortality rates for both pre-and post-implementation periods were low, because Emergency Department patients were included in the analysis for both time periods.

The MLA also fired a sepsis alert an average of two hours earlier than the SJSA, which is likely a result of the predictive design of the algorithm. The early sepsis warning provided by the MLA, coupled with its high accuracy, potentially enabled earlier clinical intervention to identify sepsis-related cases, provide supportive treatment, and possibly prevent progression of the condition. Studies have shown that early treatment of sepsis can improve patient outcomes [6, 23], and that confirmation of a positive microbiology and correspondingly targeted antibiotic therapy can improve mortality rates [24]. The early, accurate warning of sepsis onset may have provided clinicians an opportunity both to begin early treatments and to identify the causal agent in a timely manner.

The MLA’s ability to maintain currently high sensitivity and specificity, as demonstrated by its performance on retrospective data, is of clinical importance. Alarm systems with low specificity can generate high numbers of false alarms, contributing to the problem of alarm fatigue [25]. Alarm fatigue presents a patient safety concern, as providers may begin to ignore alarms which they deem unreliable. The high specificity maintained by this MLA may help mitigate alarm fatigue in clinical settings.

The algodiagnostic assessed in this study has previously been examined in several retrospective studies, where it has been validated for detection of sepsis [14], severe sepsis [13], and septic shock [15]. The algodiagnostic has also been previously evaluated in prospective studies, including a randomized controlled trial where use of the MLA resulted in statistically significant decreases in in-hospital mortality and average length of stay [17]. The present study presents further evidence that machine-learning methods for sepsis detection and prediction can provide routes towards improving sepsis-related patient outcomes.

### Limitations

The present study examines the algorithm for sepsis detection in a single medical center located in West Virginia. Other settings, with different patient demographic characteristics and EHR recording practices, may experience different outcomes with utilizing this MLA. Further, this MLA was assessed only in the emergency department and the intensive care units at CHH. The MLA may perform differently in other hospital settings, such as specialty cancer centers. The limited period of monitoring both pre-and post-implementation metrics additionally limit the generalizability of these results.

Further, confounding factors may have influenced the differences in patient outcomes noted between the pre-and post-implementation periods. Increased awareness of sepsis risk may have resulted in more rigorous bedside monitoring for sepsis during the post-implementation period. Clinicians may have more closely monitored those patients who generated an MLA alert in the post-implementation period of the study; increased attention to at-risk patients may therefore have been at least partially responsible for the improved outcomes noted.

## Conclusion

This clinical trial demonstrates improved patient outcomes through use of a machine learning-based sepsis prediction algorithm. Statistically significant reductions in the in-hospital mortality rate and hospital length of stay were obtained with this algodiagnostic, deployed concurrently with a rules-based sepsis monitoring system, over the rules-based system alone. These results are consistent with prior clinical results demonstrating improved patient outcomes with the use of machine learning-based sepsis prediction algorithms. Limitations of this study include a focus on only the emergency department and intensive care units in a single medical center, and a limited period of analysis. The MLA’s performance in a broader range of geographic regions and patient groups will be investigated in future studies.

## Acknowledgements

We gratefully acknowledge Jana Hoffman and Emily Huynh for their suggestions and help in editing this manuscript. We also acknowledge Robert Marriott and Julianna Lucci-Hensley for their insights and assistance.

## List of Abbreviations

AUROC: area under the receiver operating characteristic
CHH: Cabell Huntington Hospital
EHR: electronic health record
ICU: intensive care unit
IRB: institutional review board
LOS: length of stay
MEWS: modified early warning score
MLA: machine learning algodiagnostic
ROC: receiver operating characteristic
SE: standard error
SIRS: systemic inflammatory response syndrome
SJSA: St. John’s Sepsis Agent
qSOFA: quick sequential organ failure assessment

## Declarations

**Ethics approval and consent to participate**. Approval for the study was granted by the institutional review board (IRB) at Marshall University.

**Consent for publication**. Not applicable.

## Availability of data and materials

The data that support the findings of this study are available from CHH but restrictions apply to the availability of these data, which were used under license for the current study, and so are not publicly available. Data are however available from the authors upon reasonable request and with permission of CHH.

## Author contributions

HB, EP, DGC, and RD conceived and designed this study. All authors contributed to data analysis and interpretation. HH and ALP drafted the manuscript, and all authors revised the manuscript for critical content. All authors read and approved the manuscript.

